# Sex Differences in Effects of Obesity on Reproductive Hormones and Glucose Metabolism in Puberty: The Health Influences of Puberty Study

**DOI:** 10.1101/497339

**Authors:** Natalie Nokoff, Jessica Thurston, Allison Hilkin, Laura Pyle, Philip Zeitler, Kristen J. Nadeau, Nanette Santoro, Megan M. Kelsey

## Abstract

**Context:** Obesity is known to impact reproductive function in adults, but little is known about its effects on reproductive hormones during puberty or sex differences in these effects.

**Objective:** To assess sex differences in effects of obesity on reproductive hormones and their relationship to insulin sensitivity and secretion.

**Design:** Cross-sectional study including anthropometrics, serum and urine reproductive hormone concentrations, and intravenous glucose tolerance testing (IVGTT) to assess acute insulin response to glucose (AIR_g_) and insulin sensitivity (S_i_)

**Setting:** Outpatient academic clinical research center

**Patients:** Fifty-one normal weight (NW, BMI-Z=-0.11 ± 0.77, age=11.5 ± 1.7 years) and 53 obese (BMI-Z=2.22 ± 0.33, age=10.9 ± 1.5 years) girls (n=54) and boys (n=50), Tanner stage 2-3

**Results:** Obese boys had lower total testosterone (p<0.0001) and higher concentrations of the urinary estradiol metabolite, E1c, (p=0.046) than NW boys. Obese girls had higher free androgen index (FAI, p=0.03) than NW girls. Both obese boys and girls had lower sex hormone-binding globulin (SHBG, p<0.0001) than NW. AIR_g_ was inversely related to SHBG in boys (R^2^= 0.36, p<0.0001) and girls (R^2^=0.29, p=0.0001). Insulin resistance correlated with lower SHBG in boys (R^2^=0.45, p<0.0001) and girls (R^2^=0.24, p=0.0003), lower total testosterone for boys (R^2^=0.15, p=0.01), and higher FAI for girls (R^2^=0.08, p=0.04).

**Conclusion:** Obese youth have lower SHBG than NW youth, but obesity has differential effects on reproductive hormones in girls vs. boys, which are apparent early in puberty. Ongoing longitudinal studies will evaluate the impact of obesity on reproductive hormones in girls and boys as puberty progresses.

## Introduction

The prevalence of obesity during puberty has been rising rapidly (1). Puberty is a critical window in reproductive and metabolic health with long-lasting effects during the lifespan. Therefore, it is critical to understand the impact of obesity during pubertal development as a risk factor for reproductive and metabolic disease. In adults, obesity is associated with reproductive dysfunction, including hypogonadism in males and females and polycystic ovarian syndrome (PCOS) in females (2,3). Obesity also appears to impact pubertal development and is known to be associated with earlier puberty in girls in cross-sectional (4,5) and longitudinal studies (6). Results of studies evaluating the impact of obesity on male pubertal development are mixed, with some studies, primarily European, suggesting that a higher BMI results in earlier puberty (7–9) while others, primarily American, suggest later pubertal development (10–12). Results of a limited number of studies, largely cross-sectional, assessing differences in serum testosterone in obese vs. normal weight (NW) boys have been mixed, with some studies, including several from our group, showing lower total testosterone in obese boys (13–16) and others showing no difference (17). In girls, there is an association between obesity and hyperandrogenemia (18,19). In addition to lack of clarity about the immediate impact of obesity on sex hormones in youth, the timing of when differences in sex hormones in obese youth begin to reflect patterns seen in adults is unclear.

In NW adolescents, there is a physiologic decrease in insulin sensitivity during puberty that begins at Tanner Stage 2 (T2), reaches a nadir at T3, and returns to near prepubertal levels sometime after puberty is complete (20–22). This pubertal insulin resistance is thought to play an important role in the pathophysiology of youth-onset type 2 diabetes mellitus (T2DM) in obese youth, which is a disease that rarely presents prior to puberty (23). Youth-onset T2DM is twice as common in girls as in boys (24), a sex difference not seen in adults. The reasons for this sex difference are unknown and there are significant additional knowledge gaps in how obesity impacts the relationship between reproductive hormones and glucose metabolism in girls and boys, particularly in early puberty. Most studies assessing the effects of obesity on sex steroids, insulin sensitivity, and β-cell function are cross-sectional (16, 18,25) and very few involve youth in early puberty. To address these gaps, studies using rigorous measures of insulin sensitivity, β-cell function and reproductive hormones, and including all stages of puberty are needed.

Our primary objective was to assess sex differences in the relationship between obesity and reproductive hormones in early puberty. We hypothesized that there would be differences in sex steroid patterns between obese and NW youth in early puberty, similar to patterns in adults, with higher bioavailable androgens in obese than NW girls and lower testosterone in obese than NW boys. Given the known relationships among insulin sensitivity, insulin secretion and sex steroids in adults, we also conducted exploratory analyses to examine associations among insulin sensitivity (S_i_), insulin secretion (acute insulin response to glucose, AIR_g_) and sex steroids in early pubertal youth.

## Methods

### Participants

NW and obese girls and boys in early puberty (T2-3) were recruited between 2009 and 2015 for the Health Influences of Puberty (HIP) Study, a longitudinal study evaluating the effects of obesity on glucose metabolism during puberty. Recruitment took place in the primary care and multidisciplinary obesity clinics at Children’s Hospital Colorado, as well as from local pediatric clinics through advertisements. The study was approved by the Colorado Multiple Institutional Review Board and consent and assent were obtained from all participants.

NW (BMI 5^th^-85^th^ percentile) and obese (BMI ≥ 95^th^ percentile) youth, who were T2-3 and age ≥ 9 years at the time of enrollment, were included. All obese youth were screened with an oral glucose tolerance test (1.75 g/kg of glucola [maximum 75g]); participants with impaired fasting glucose ([IFG], fasting glucose 100-125 mg/dl), impaired glucose tolerance ([IGT], 2-hr glucose 140-199 mg/dl), or T2DM (fasting glucose ≥ 126 mg/dl, 2 hr. glucose ≥ 200 mg/dl) using ADA criteria (26) were excluded. Exclusion criteria for all participants also included: specific genetic syndromes or disorders known to affect glucose tolerance other than diabetes; oral steroids within the last 60 days or high-dose inhaled (>1000 mcg daily) steroids; medication(s) that are known to cause weight gain or weight loss, including atypical antipsychotics; insulin sensitizer use within the last year; use of anabolic steroids; hypertension or hyperlipidemia requiring pharmacological intervention; proteinuria, weight >300 pounds (due to dual-energy X-ray absorptiometry [DXA] limits) or other significant organ system illness or condition (including psychiatric or developmental disorder) that would prevent full participation. NW participants were not screened for IGT, IFG or T2DM, but were excluded for a known diabetes diagnosis.

The screening visit also included pubertal staging, performed by a pediatric endocrinologist. Puberty was assessed using the standards of Tanner and Marshall for breast development in girls (using inspection and palpation), genital development in boys, and pubic hair in boys and girls (27,28). Because of the subjectivity of Tanner genital staging for boys, testicular volume was also assessed using a Prader orchidometer and assigned a Tanner stage equivalent as follows: T1 < 4 mL; T2 ≥ 4 mL and < 8; T3 ≥ 8 and < 12 mL; T4 ≥ 12 and ≤ 15 mL; T5 >15 mL. Girls and boys needed to be ≥ T2 for breast development or testicular volume, respectively, and no more than T3 for breast development, testicular volume, or pubic hair to be eligible for study entry. Breast development and testicular volume took priority over pubic hair for assignment of Tanner stage.

### Research visit

Approximately 3 weeks after the screening visit, participants were admitted to the Children’s Hospital Colorado Clinical and Translational Research Center (CTRC) following a 3-day standard macronutrient diet (55% carbohydrates, 30% fat, and 15% protein) provided by the metabolic kitchen and following 3 days of restriction of vigorous physical activity. Participants fasted overnight prior to admission. Height was measured on a Harpenden stadiometer and weight on a digital electronic scale. Height and weight were recorded to the nearest 0.1 centimeter and kilogram, respectively. Serum and plasma were drawn on a morning fasting sample for the following assays: glucose, insulin, dehydroepiandrosterone sulfate (DHEA-S), estradiol, total testosterone, and SHBG.

Participants then underwent a frequently sampled intravenous glucose tolerance test (IVGTT): participants had an intravenous (IV) catheter inserted into each arm. Baseline blood samples were drawn for fasting plasma glucose and serum insulin. Participants then had an infusion of 0.3 g/kg of 25% dextrose given over 90 seconds. Glucose and insulin concentrations were sampled from the IV in the contralateral arm at times 2, 3, 4, 5, 6, 8, 10, 12, 14, 16, 19, 22, 25, 30, 35, 40, 50, 60, 70, 80, 90, 100, 120, 140, 160 and 180 minutes. Bergman’s minimal model program was used to calculate S_i_, AIR_g_, and the disposition index (DI) (29). DXA was used to measure percent fat mass.

### Laboratory assays

Laboratory assays were performed by the University of Colorado Anschutz CTRC core laboratory. Glucose was measured by enzymatic UV testing (AU480 Chemistry Analyzer, Beckman/Coulter, Brea, CA), inter- and intra-assay coefficient of variation (CV) 1.44% and 0. 67%, respectively and sensitivity 10 mg/dL. Insulin was measured by radioimmunoassay (EMD Millipore, Darmstadt, Germany), inter- and intra-assay CV 9.8% and 5.2%, respectively and sensitivity 3 uIU/mL. DHEA-S was measured by chemiluminescence (Beckman Coulter, Brea, CA), inter- and intra-assay CV 3.4% and 2.3%. respectively and sensitivity 2 ug/dL. Estradiol was measured by chemiluminescence (Beckman Coulter, Brea, CA), inter- and intra-assay CV 8.2% and 4.3%, respectively and sensitivity 10.0 pg/mL. SHBG was measured by chemiluminescence (Beckman Coulter, Brea, CA), inter- and intra-assay CV 5.7% and 3.6%, respectively and sensitivity 3 nmol/L. Total testosterone was measured by radioimmunoassay of serum (Beckman Coulter, Brea, CA), inter- and intra-assay CV 5.1% and 2.1%, respectively and sensitivity 17 ng/dL. Free androgen index (FAI) was calculated as the ratio of total testosterone to SHBG ([testosterone/SHBG]*100).

Urine studies: follicle-stimulating hormone (FSH), luteinizing hormone (LH) and estrone metabolites (E1c, a mixture of estrone sulfate and estrone glucuronide) were measured by DELFIA immunoflourometric assay platform (in-house ELISA) on first morning urine sample and normalized to creatinine (30,31). FSH (mLU/mgCr), inter- and intra-assay CV 6.6% and 5.0%, respectively and sensitivity 0.98 IU/L. LH (mLU/mgCr), inter- and intra-assay CV: 4.8% and 3.4%, respectively, and sensitivity 0.57 IU/L. E1c (ng/mgCr), inter- and intra-assay CV 6.8 and 2.2%, respectively, and sensitivity 1 pg/well (32).

### Statistical methods

Descriptive statistics were calculated and groups (NW vs. obese) were compared using linear regression models adjusted for pubertal stage. Variables were assessed for normality and, after evaluating diagnostics, AIR_g_ and Si were log-transformed. Univariable linear regression models were utilized to examine the associations between outcomes (Si, AIR_g_, DI and BMI-Z) and predictors (E1c, total testosterone, FAI, LH, FSH, and SHBG). Pearson correlations were utilized to examine associations between BMI-Z and Si, AIR_g_ and DI. Due to cohort differences in racial/ethnic composition in NW and obese youth, all analyses were performed with and without adjustment for race/ethnicity. Analyses were conducted using SAS version 9.4 (SAS Institute, Cary, NC). BMI-Z was calculated using the US Centers for Disease Control and Prevention SAS code. This was a secondary analysis of effects of obesity on reproductive hormones in a study that was primarily designed to evaluate effects of obesity on glucose metabolism during puberty, and therefore no adjustments were made for multiple testing.

## Results

Demographic characteristics by group are in Table 1. Fifty boys (27 NW, 23 obese, average age 12.3 + 1.4 years) and 54 girls (24 NW, 30 obese, average age 10.2 + 1.0 years) completed the study visit. Obese and NW youth were similar for age and Tanner stage distribution within each sex. There were significant differences for race among boys and ethnicity among girls between the NW and obese cohorts, with more minority representation in the obese participants.

**Table 1:** Baseline demographics and laboratory values for normal weight (NW) and obese boys and girls

| Variable | Girls (n=54) |  | p value | Boys (n=50) |  | p value |
| --- | --- | --- | --- | --- | --- | --- |
|  | NW (n=24)<br>Median<br>(IQ range) | Obese (n=30)<br>Median<br>(IQ range) |  | NW (n=27)<br>Median<br>(IQ range) | Obese (n=23)<br>Median<br>(IQ range) |  |
| Age (years) | 10 (10, 11) | 10 (9, 11) | 0.13 | 12 (12, 14) | 12 (11, 13) | 0.23 |
| Tanner stage n (%) | T2: 14 (58.3)<br>T3: 10 (41.7) | T2: 17 (56.7)<br>T3: 13 (43.3) | 0.90 | T2: 18 (66.7)<br>T3: 9 (33.3) | T2: 17 (73.9)<br>T3: 6 (26.1) | 0.58 |
| Race n (%) | 14 (58.3) | 22 (73.3) | 0.23 | 18 (66.7) | 18 (78.3) | 0.0005 |
| White | 7 (29.2) | 3 (10.0) |  | 3 (11.1) | 3 (13.0) |  |
| Black | 3 (12.5) | 5 (16.7) |  | 6 (22.2) | 2 (8.7) |  |
| Other |  |  |  |  |  |  |
| Ethnicity n (%) |  |  | 0.01 |  |  | 0.44 |
| Hispanic | 10 (41.7) | 23 (76.7) |  | 5 (18.5) | 16 (69.6) |  |
| Non-Hispanic | 12 (50.0) | 7 (23.3) |  | 21 (77.8) | 7 (30.4) |  |
| Unknown | 2 (8.3) | 0 |  | 1 (3.7) | 0 |  |
| BMI %ile | 48.4 (27.6, 65.3) | 98.1 (97.2, 99.3) | **<0.0001** | 47.3 (23.8, 67.7) | 98.8 (98.5, 99.5) | **<0.0001** |
| BMI Z-score | -0.04 (-0.6, 0.4) | 2.08 (1.9, 2.5) | **<0.0001** | -0.1 (-0.7, 0.5) | 2.3 (2.2, 2.6) | **<0.0001** |
| Body fat (%) | 27.2 (22.9, 32.2) | 44.3 (41.6, 47.0) | **<0.0001** | 23.9 (17.2, 29.2) | 43.1 (36.4, 45.5) | **<0.0001** |
| Serum estradiol (pg/mL) | 17 (5, 27.5) | 11.5 (5, 24) | 0.14 | 10 (5, 18) | 10 (6.9) | 0.93 |
| Corrected* urinary FSH (mLU/mgCr)^ | 9.4 (2.4) | 0.2 (2.8) | **0.01** | 4.7 (6.7) | 3.9 (0.4, 4.0) | 0.51 |
| Corrected* urinary LH (mLU/mgCr)^ | 2.0 (0.9) | 0.01 (1.1) | 0.17 | 0.7 (0.8) | 0.8 (0.9) | 0.94 |
| DHEA-S (µg/dL) | 48 (28, 90.5) | 60 (45, 89) | 0.69 | 101 (61, 153) | 140 (96, 190) | 0.1 |
| Insulin sensitivity (S<sub>i</sub>) (x10<sup>-4</sup>/min-1/mIU/mL) | 5.9 (3.9, 8.3) | 2.2 (1.5, 3.0) | **0.013** | 8.69 (4.9, 12.1) | 2.0 (1.5, 3.0) | **<0.0001** |
| Insulin secretion (AIR<sub>g</sub>) (µIU/mL) | 714 (314, 792) | 1923 (962, 2664) | **<0.0001** | 403 (324, 629) | 1805 (1032, 3356) | **<0.0001** |
| Disposition Index (DI) (x10<sup>-4</sup>/min-1) | 3749 (2869, 4750) | 4139 (2988, 4774) | 0.35 | 3034 (2633, 5239) | 4088 (2868, 5354) | 0.62 |

\*Normalized to creatinine. BMI = body mass index, FSH = follicle stimulating hormone, LH = luteinizing hormone, DHEA-S = dehydroepiandrosterone sulfate. ^Least squared mean (standard error); anthropometric, hormonal and metabolic comparisons are adjusted for Tanner stage and race/ethnicity

### Relationships of obesity to reproductive hormones and SHBG

Baseline hormone concentrations are shown in Table 1. As shown in Figure 1, obese boys had significantly lower SHBG (median, [IQ range], 18 [14, 24] nmol/L vs. 57 [47, 73], p<0.0001), lower total testosterone (47 [23, 90] ng/dL vs. 81 [48, 296], p<0.0001) and higher urinary E1c (11.3 [6.7, 26.5] ng/mgCr vs. 6.2 [0.9, 7.4], p=0.046) than NW boys, but there was no significant difference in FAI (213.6 [156.3, 360.0] vs. 164.8 [64.8, 416.9], p=0.9).

**Figure 1:**
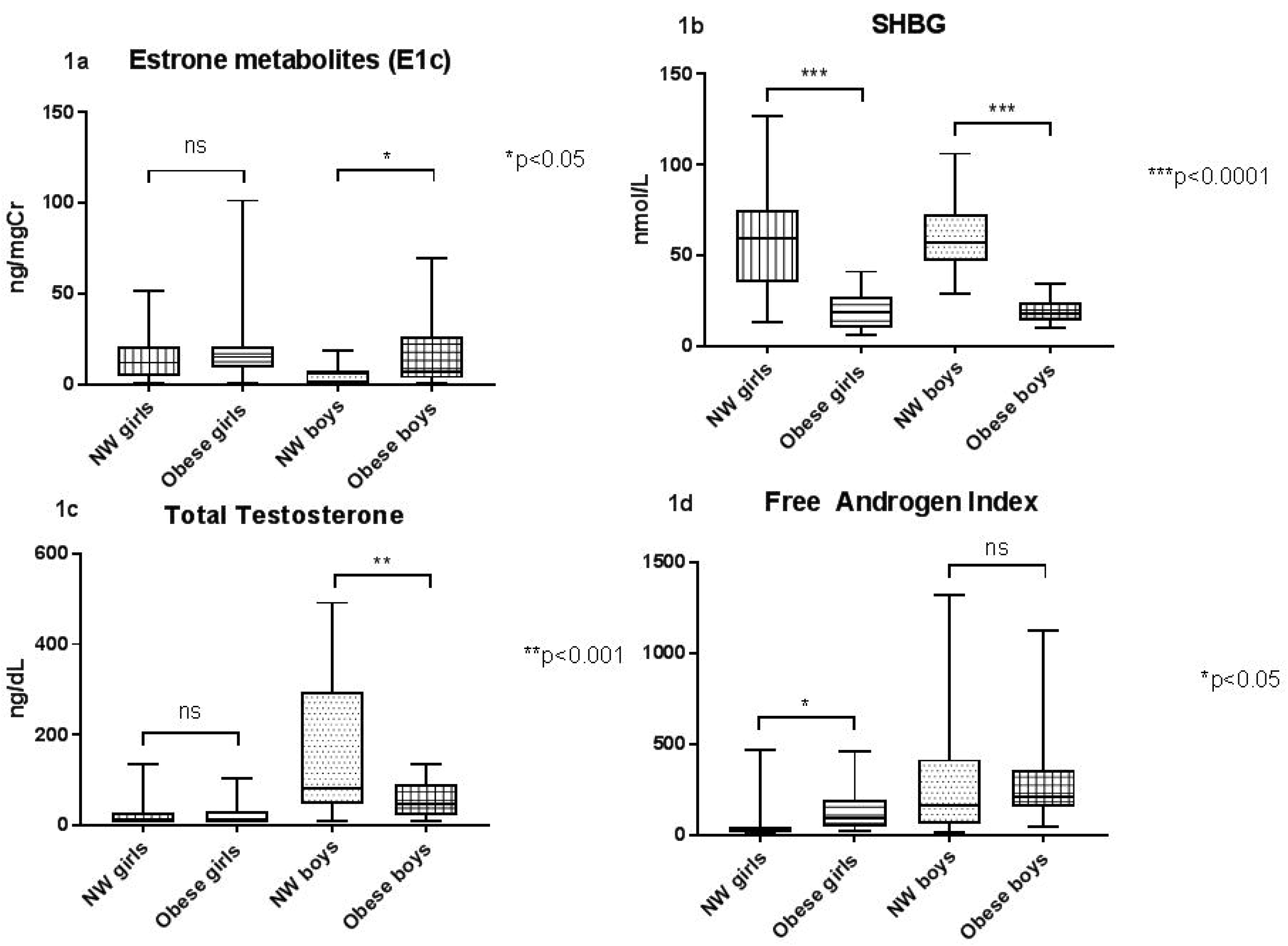
The box plots show the minimum, 25^th^ percentile, median, 75^th^ percentile and maximum values for: estradiol metabolites (E1c) in 1a, sex hormone-binding globulin (SHBG) in 1b, total testosterone in 1c, and free androgen index (FAI) in 1d for normal weight (NW) and obese girls and boys; ns = not significant.

Obese girls also had significantly lower SHBG (18.5 [10, 27] nmol/L vs. 59.5 [35.5, 75], p<0.0001), but had higher FAI (91.7 [44.7, 195] vs. 26.7 [13.5, 44.9], p=0.03) than NW girls. There were no significant differences in E1c (15.2 [9.3, 43.8] ng/mgCr vs. 12.1 [4.9, 21.1], p=0.18) or total testosterone (8.5 [8.5, 31] ng/dL vs. 8.5 [8.5, 28.5], p=0.77, Figure 1) between obese and NW girls, adjusted for Tanner stage. Obese girls had a lower urinary FSH than NW girls (2.8 [1.3, 7.0] vs. 3.6 [2.8, 14.2] mU/mgCr) in analyses adjusted for baseline Tanner stage and race/ethnicity (p=0.01) and for race/ethnicity alone (p=0.02), but not in unadjusted analysis (p=0.17). There were no other significant differences in urinary gonadotropin concentrations between NW and obese boys or girls (Table 1).

### Relationship between glucose metabolism and reproductive hormones

Baseline measures of insulin sensitivity and secretion are shown in Table 1. As expected, both obese boys and girls had significantly lower S_i_ and higher AIR_g_ than NW boys and girls (Table 1). There was a higher AIR_g_ was associated with higher BMI-Z in both boys (R=0.74, p<0.0001) and girls (R=0.62, p<0.0001) and a lower S_i_ was associated with higher BMI-Z in both boys (R= −0.73, p<0.0001) and girls (R= −0.54, p<0.0001).

In boys (Table 2), higher AIR_g_ was associated with higher DHEA-S and LH and lower SHBG and total testosterone. Inversely, as expected, higher S_i_ was associated with higher SHBG and total testosterone and lower DHEA-S, LH and FSH. The relationship between FSH and AIR_g_ was no longer significant (p=0.07) after adjustment for race/ethnicity. Although urinary E1c was higher in the obese boys compared to NW boys, BMI-Z did not correlate with E1c (R^2^= 0.11, p=0.08). In girls (Table 2), higher AIR_g_ and was associated with higher FAI and lower SHBG. Conversely, higher S_i_ was associated with higher SHBG and lower FAI.

**Table 2:** Univariate regression analyses for boys and girls between outcomes (S_i_, AIR_g_) and predictors (total testosterone, FAI, DHEA-S, SHBG)

| Dependent variable (log-transformed) | Parameter | Estimate | $R^2$ | p value |
| --- | --- | --- | --- | --- |
| **Boys** |  |  |  |  |
| $AIR_g$ | SHBG | -0.025 | 0.36 | <.0001 |
| $AIR_g$ | DHEA-S | 0.008 | 0.19 | 0.003 |
| $AIR_g$ | Total testosterone | -0.004 | 0.22 | 0.001 |
| $AIR_g$ | FSH | 0.138 | 0.17 | 0.04 |
| $S_i$ | SHBG | 0.024 | 0.45 | <.0001 |
| $S_i$ | DHEA-S | -0.006 | 0.18 | 0.004 |
| $S_i$ | Total testosterone | 0.003 | 0.15 | 0.01 |
| $S_i$ | FSH | -0.166 | 0.26 | 0.008 |
| $S_i$ | LH | -0.368 | 0.22 | 0.01 |
| **Girls** |  |  |  |  |
| $AIR_g$ | SHBG | -0.017 | 0.29 | <.0001 |
| $AIR_g$ | FAI | 0.003 | 0.13 | 0.009 |
| $S_i$ | SHBG | 0.016 | 0.24 | <0.001 |
| $S_i$ | FAI | -0.002 | 0.08 | 0.042 |

Significant results are shown. $S_i$ = insulin sensitivity, $AIR_g$ = acute insulin response to glucose, FAI = free androgen index, DHEA-S = dehydroepiandrosterone sulfate, SHBG = sex hormone-binding globulin.

## Discussion

The HIP study is one of the few studies that explores the impact of obesity on reproductive hormones in early puberty and one of the first to evaluate interactions between rigorous estimates of insulin sensitivity and secretion and reproductive hormones during this critical developmental period. In addition to providing insight into the biological effects of obesity during this critical window for long-term reproductive function, it is important to understand the contribution of sex hormones in obese youth to the pathophysiology of youth-onset diabetes, the onset of which is closely associated with puberty, and other puberty-associated metabolic abnormalities. Studies attempting to intervene during later puberty to improve diabetes outcomes in obese youth with pre-diabetes and T2DM have been unsuccessful (33,34); thus intervention may need to occur earlier in puberty. The HIP study is also novel in employing measurement of gonadotropins and estradiol metabolites from overnight urine samples. This is a more sensitive approach to measuring low concentrations of sex hormones than those previously employed, such as random or first morning serum measurements, which is critical for examination of early puberty.

While both obese boys and girls have much lower SHBG, an independent predictor of type 2 diabetes incidence in adults (35), than NW youth, obesity appears to have sex-specific effects on sex hormone concentrations early in puberty. Early pubertal obese girls have a higher FAI but no differences in total testosterone or urinary E1c compared to NW girls. In contrast, obese boys have higher E1c and lower total testosterone, but no difference in FAI compared to NW boys. Measurement of total testosterone is affected by SHBG, as 30-44% of total testosterone is tightly bound to SHBG (36,37), whereas it has less of an effect on estradiol measurement. Biologically active or “bioavailable” testosterone is thought to represent a combination of free testosterone and that which is loosely bound to albumin. Thus, in the HIP Study, the obesity effects on testosterone concentrations are likely driven by lower SHBG, resulting in higher FAI in girls and lower total testosterone in boys. It is important to note that FAI (a measure of bioavailable testosterone) was no different between NW and obese boys. This demonstrates the importance of measuring total and free testosterone when assessing gonadal function in obese boys. This study also adds to the literature by demonstrating that these hormonal associations with obesity are apparent very early on in puberty, confirming findings from other studies focusing on youth in later puberty.

While bioavailable testosterone was not affected by obesity in these pubertal boys, higher gonadotropins and lower testosterone were associated with lower Si, potentially suggesting that insulin resistance could be associated with a primary hypogonadism (i.e., low testosterone, elevated gonadotropins) early in puberty. This is opposite to what is reported in obese adult men, who are presumed to have hypogonadotropic hypogonadism (38); however, there is still some controversy as to the true cause of hypogonadism and our results from early puberty suggest that, at least initially, insulin resistance may directly affect the testis. Thus, though the associations between low total testosterone and low S_i_ and high AIR_g_ in early pubertal boys are more likely a reflection of the impact of hyperinsulinemia on SHBG, the low testosterone may be resulting in a drive to increase gonadotropin secretion from the pituitary.

In addition to impacting measurement of sex steroids, low SHBG is thought to have cardiometabolic implications. In the HIP study, the markedly lower SHBG concentrations in early pubertal obese youth were directly associated with S_i_ and inversely associated with AIR_g_, consistent with previous reports in adults and later pubertal youth (13, 16, 17). SHBG is known to be associated with insulin resistance, likely due to direct suppression of SHBG by compensatory hyperinsulinemia (39), and has been demonstrated to be an independent predictor of diabetes in adults, after adjusting for a number of factors (age, cardiovascular disease, smoking, alcohol, socioeconomic status, BMI, and waist circumference) (40), but little is known about SHBG as a risk marker for cardiometabolic disease in youth. Longitudinal studies are now needed to determine whether SHBG in early puberty youth is a useful cardiometabolic disease risk marker of later disease progression.

Our findings are similar to several other studies showing lower total testosterone in obese pubertal boys (13–15) and demonstrate that these differences are detectable as early as Tanner 2, when total testosterone is present in low concentrations. It is important to note that a radioimmunoassay (RAI) was used in this study, which may have decreased sensitivity at the low testosterone concentrations seen in women of all ages and in early pubertal boys, and therefore total testosterone measurement by liquid chromatography-tandem mass spectrometry (LC-MS/MS) is preferred in these groups (41). Studies that have utilized LC-MS/MS have shown that prepubertal obese and NW boys have similar total testosterone (16), but that total testosterone becomes lower in obese boys as they progress through puberty (16,42). The low total testosterone but normal FAI in our and other cohorts of obese pubertal boys has important clinical implications, suggesting the necessity of measuring free testosterone rather than total testosterone when assessing gonadal dysfunction in obese boys. It remains unclear however, when the impact of obesity on free testosterone, seen in adult males (43), emerges.

Among obese girls in the HIP Study, the exposure FAI was about three times as high in obese pubertal girls compared to NW girls of the same pubertal stage, reflecting early evidence of obesity-associated hyperandrogenism (3), again demonstrating the impact of obesity on reproductive hormones early in puberty and confirming previous findings in later pubertal girls (17). Unlike other studies that have found associations between high BMI and DHEA-S (14, 17,18), but had a wider range in pubertal stages, we did not find differences in DHEA-S in our early pubertal cohort. However, we did find a weak association between DHEA-S and Si and AIR_g_ in the HIP girls, indicating that the previous associations may be more related to insulin resistance than to the obesity itself. Again, longitudinal studies using sensitive LC-MS/MS measurement of androgens are now needed to better understand the impact of obesity on sex steroids and adrenal androgens as puberty progresses in girls.

The use of a first morning urine sample measure gonadotropins and E1c in the HIP Study had several advantages. Sex steroids and gonadotropins are primarily secreted at night, in a pulsatile manner and in low concentrations in early puberty (44). Measuring gonadotropins and E1c on a first morning urine collection (normalized to creatinine) reflects the overnight hormone production yet avoids frequent overnight blood sampling. Urinary reproductive hormone assays have been validated in adults (45,46) and have also been studied in children (47,48). In adult women, urine and serum concentrations of estradiol and E1c are highly correlated (R=0.93) (46). Furthermore, urinary LH, FSH, E1c, and progesterone metabolites have been show to follow the expected pattern based on serum values in a cohort of girls ages 11-13 years followed prospectively over 2 years (48), validating the use of these assays in girls in early puberty. There have been no prior studies of urinary E1c in boys in early puberty. Using this sensitive assay, we have been able to demonstrate, for the first time, the presence of higher estrogen exposure in obese than lean boys at the earliest stage of puberty. It has been proposed that increased aromatization of androgens in adipose (49) leads to higher estradiol concentrations in obese men, potentially driving the higher incidence of hypogonadotropic hypogonadism in these men through negative feedback of estradiol on the pituitary and hypothalamus. The presence of obesity and high estradiol going into puberty may, therefore, further impact reproductive function through negative feedback on the pituitary as puberty progresses, in addition to the insulin resistance-related primary hypogonadism mentioned above.

In conclusion, our study demonstrates reproductive implications of the presence of obesity at the onset of puberty and illustrates sex differences in this impact. Importantly, this study demonstrates that some of the sex steroid patterns seen in obese adults, including higher E1c and lower total testosterone in obese men and higher bioavailable testosterone in obese women, are already present in early pubertal obese youth. In contrast, the lower bioavailable testosterone seen in obese adult males was not present in our early pubertal obese boys. The timing of the emergence of the hypoandrogenemia reported in obese adult males remains unclear. Further longitudinal study is needed to better understand the impact of these early changes in obese youth on long-term reproductive and cardiometabolic function.

## Acknowledgements

The authors wish to thank the participants in the study. This research was supported by Dr. Kelsey’s American Diabetes Association Junior Faculty Award (1-11-JF-23) and the Children’s Hospital Colorado Research Institute Research Scholar Award. Drs. Kelsey and Santoro are supported by the Colorado Interdisciplinary Research Careers in Women’s Health NIH/NICHD BIRCWH K12 (HD057022-06). Institutional funding: NIH/NCATS Colorado CTSA UL1 TR001082. Dr. Nokoff is supported by a T32 grant (T32 DK 63687).

